# YY1 is a transcriptional activator of mouse LINE-1 Tf subfamily

**DOI:** 10.1101/2024.01.03.573552

**Authors:** Karabi Saha, Grace I. Nielsen, Raj Nandani, Lingqi Kong, Ping Ye, Wenfeng An

## Abstract

Long interspersed element type 1 (LINE-1, L1) is an active autonomous transposable element (TE) in the human genome. The first step of L1 replication is transcription, which is controlled by an internal RNA polymerase II promoter in the 5’ untranslated region (UTR) of a full-length L1. It has been shown that transcription factor YY1 binds to a conserved sequence motif at the 5’ end of the human L1 5’UTR and dictates where transcription initiates but not the level of transcription. Putative YY1-binding motifs have been predicted in the 5’UTRs of two distinct mouse L1 subfamilies, Tf and Gf. Using site-directed mutagenesis, in vitro binding, and gene knockdown assays, we experimentally tested the role of YY1 in mouse L1 transcription. Our results indicate that Tf, but not Gf subfamily, harbors functional YY1-binding sites in its 5’UTR monomers. In contrast to its role in human L1, YY1 functions as a transcriptional activator for the mouse Tf subfamily. Furthermore, YY1-binding motifs are solely responsible for the synergistic interaction between monomers, consistent with a model wherein distant monomers act as enhancers for mouse L1 transcription. The abundance of YY1-binding sites in Tf elements also raise important implications for gene regulation at the genomic level.

## Introduction

Transposable elements (TEs) constitute at least 46% of the human genome (1). The vast majority of human TEs are retrotransposons, which replicate in the genome through an RNA intermediate and are further divided into four classes: long terminal repeat (LTR) element, long interspersed element (LINE), short interspersed element (SINE), and the composite SVA element (1). LINE-1 (L1) is of particular interest as it is the only class of TEs that are both autonomous and active in the human genome (2–5). The vast majority of L1s in the human genome are 5’ truncated. A full-length L1 consists of a 5’ untranslated region (5’UTR), two tandem open reading frames (ORF1 and ORF2) that are separated by a short inter-orf spacer, and a 3’ untranslated region (3’UTR) (6). The transcription of a full-length L1 mRNA is controlled by an internal Pol II promoter in the 5’UTR (7,8). L1 mRNA has two essential functions during L1 retrotransposition: first, being translated into ORF1 and ORF2 proteins (9,10); second, being reverse transcribed by ORF2 protein into a new DNA copy (11,12). Thus, controlling L1 promoter activity represents a critical regulatory step for L1 replication cycle (13,14). One key aspect of L1 transcriptional regulation involves epigenetic mechanisms, such as DNA methylation, histone modification and, in germ cells, Piwi-interacting RNAs (piRNAs) (15). Another important layer of L1 transcriptional regulation is the availability and binding of transcriptional factors to the L1 promoter region (14).

Transcription factor YY1 is a member of the C2H2 zinc finger protein family and is evolutionarily conserved among animals (16). In fact, human and mouse YY1 proteins are 98.6% identical to each other over the length of 414 amino acids (17,18). It is ubiquitously expressed in cell lines and human tissues (18–21). Since its initial discovery, YY1-binding sites have been found in promoters of many cellular and viral genes, activating or repressing them in a context dependent manner (22). Several models have been proposed to explain YY1’s seemingly divergent roles, including its recruitment of coactivators or corepressors (22). Recent studies indicate that YY1 is a structural regulator of enhancer-promoter interactions (21,23). It occupies active enhancers and promoter-proximal sequences across cell types (21,23). Through dimerization YY1 mediates interactions between enhancers and promoters at a genome-wide scale (21). Its association with these regulatory sequences is dependent on a known DNA-binding motif and augmented by YY1’s binding to RNA (21). Downregulation or ablation of YY1 leads to widespread changes in gene expression, with similar number of genes upregulated or downregulated (21,24), suggesting a model in which YY1 coordinates the positioning of other activators or repressors at individual promoters (21).

YY1 plays an important role in human L1 transcription. Human L1 5’UTR harbors a consensus YY1-binding site at nucleotide position (nt) 9-20 (25). The binding site only has marginal effect for the overall transcriptional output from the full-length 5’UTR (26) despite being critical for the activity of the first 155 bp of L1 5’UTR (25). Instead, an intact YY1-binding motif controls the transcription initiation from the 5’ end of the 5’UTR (26). However, whether YY1 regulates mouse L1 transcription has been inconclusive. Twenty-nine L1 subfamilies have amplified in the mouse genome since the split between mouse and rat about 13 million years ago (27). In the last one million years, at least four mouse L1 subfamilies (A_I, Tf_I, Tf_II, and Gf_I) have been active, with the average age of elements within each subfamily varying from 0.21, 0.25, 0.27, to 0.75 million years, respectively (27). Like human L1, mouse L1 5’UTRs possess promoter activities (28–31). In contrast to human L1, mouse 5’UTRs are organized into tandemly repeated monomers (27,32,33). Recently, we have shown that two-monomer consensus sequences from six L1 subfamilies differ in their promoter activities when tested in two separate murine cell lines in reporter assays (34). Putative YY1-binding motifs have been predicted in both Tf (30,31,35) and Gf monomers (36). A previous study attempted to determine the function of the putative YY1-binding site in Tf 5’UTRs by comparing the promoter activity of three genomic Tf promoters: one with a YY1 site and two without (37). Although a two-fold reduction in promoter activity was observed for the latter two Tf promoters these results are inconclusive due to the presence of other confounding mutations.

In this study we aim to determine the function of putative YY1-binding sites in the 5’UTR of mouse Tf_I and Gf_I subfamilies by performing site-directed mutagenesis, in vitro binding assays, and siRNA knockdown of YY1 protein in two murine cell lines. We found that the putative YY1-binding site is functional and required for Tf promoter while the predicted YY1-binding site in Gf_I promoter is not functional due to one nucleotide substitution in the core binding motif. Consistent with a regulatory role of YY1 for Tf transcription, our analysis of YY1 ChIP-seq data showed that YY1 occupancy is enriched at Tf_I/II but not at other evolutionarily young L1 subfamilies in mouse embryonic stem cells.

## Materials and Methods

### Plasmid construction

A detailed list of the promoter constructs, including the corresponding promoter sequences, is provided as a supplemental table (Table S1). In all promoter constructs, the respective L1 promoter variant is positioned immediately upstream of the firefly (Fluc) reporter gene and flanked by two heterotypic SfiI sites (SfiI_L=GGCCAAAA/TGGCC and SfiI_R=GGCCTGTC/AGGCC; “/” indicates the cleavage site). The double-SfiI cassette enables directional insert swapping via a single, robust restriction/ligation cycle (38). pCH117 is a positive control vector that contains the human L1RP 5’UTR as the “L1 promoter” (34). pLK037 is a negative control vector that contains an empty double-SfiI cassette upstream of the Fluc reporter gene (34). Wild-type Tf_I promoter constructs (M2/pLK057, M1/pLK056, M1-T/pLK047, and M2-M1-T/pLK050) and wild-type Gf_I promoter constructs (M2/pLK063, M1/pLK062, and M2-M1-T/pLK051) have been described previously (34). Promoter variants containing nucleotide substitutions at the predicted YY1-binding motif were ordered as synthetic DNA fragments flanked by SfiI_L and Sfil_R restriction sites from either Genewiz (part of Azenta Life Sciences) or Twist Biosciences. Each synthetic DNA fragment was digested by SfiI (New England Biolabs) and ligated into SfiI digested backbone from pCH117 using T4 DNA ligase (New England Biolabs). All promoter variants were verified by Sanger sequencing (Elim Biopharmaceutics Inc).

### Dual-luciferase promoter assay

F9 mouse embryonal carcinoma cell line (ATCC CRL-1720) was gifted by Dr. Michael Griswold, Washington State University. A subline of NIH/3T3 mouse embryonic fibroblast cells (ATCC CRL-1658) was maintained in our lab and has been recently authenticated (34). Both cell lines were propagated in a complete culture medium composed of DMEM/High Glucose, 1% SG-200, and 10% fetal bovine serum (all from Cytiva Life Sciences). A reverse transfection protocol using Lipofectamine 3000 (Invitrogen) was followed (34). For F9 cells, a 96-well plate was coated with 0.1% gelatin for at least 30 minutes before adding transfection mix and cell suspension. Four replicate wells were allocated for each plasmid. In each well, 5 µL of transfection mix containing 10 ng of plasmid DNA was added followed by 100 µL of cell suspension (40,000 cells). The plate was incubated in a CO_2_ incubator at 37°C for 24 hours before luminescence readout. For NIH/3T3 cells, 20,000 cells were added per well, and plates were incubated for 48 hours. Dual-Luciferase Reporter Assay System (Promega) was used to measure luciferase activities. Briefly, cells were lysed using 1x passive lysis buffer and transferred to a solid white flat-bottom 96-well plate (Greiner Bio-One). Firefly luciferase activity was read first on a GloMax Multi Detection System (Promega) followed by the measurement for Renilla luciferase activity. Signal integration time was set to one second per well. Mock transfected cells and empty wells were included to evaluate the assay background.

Promoter assays under siRNA knockdown of YY1 protein were performed in F9 and NIH/3T3 cells with modifications. A mouse Yy1-specific siRNA or the AllStars Negative Control siRNA (Qiagen) was co-transfected with an L1 promoter reporter plasmid using Lipofectamine 3000 (Invitrogen). The target sequences for mouse Yy1-specific siRNAs are: Mm_Yy1_1 CACATCTTAACACACGCTAAA, Mm_Yy1_5 CAGGAGTGTGATTGGGAATAA, Mm_Yy1_6 CAGAAATTGGAAGCAAATAAA, Mm_Yy1_7 CACAAAGATGTTCAGGGATAA. The siRNA transfection mix was prepared separately without P3000 and added to the 96-well plate after the addition of plasmid DNA transfection mix. A no siRNA control was included to evaluate the effect of RNA cotransfection on the promoter assay. Per well 5 pmol siRNA was used, equivalent to a concentration of 50 nM in 100 µL culture medium. After the addition of transfection mixes, 20,000 F9 cells or 10,000 NIH/3T3 cells were added to the well. Transfected F9 cells and NIH/3T3 cells were incubated for 48 hours and 72 hours, respectively, before luciferase measurements. Four replicate wells were allocated for each condition.

### Western blot

To verify siRNA knockdown efficiency, transfection was scaled up from a 96-well to 24-well plate by a factor of 6. siRNA transfection was set up in a similar way as the dual-luciferase assay in 96-well plate. Four different mouse Yy1-specific siRNAs were transfected separately as well as in combination. A total of 30 pmol siRNA was added per well. AllStars was used as a negative control. 60,000 NIH/3T3 cells were added to each well. After 72 hours of incubation, whole cell lysate was prepared using RIPA buffer (Nalgene) in the presence of protease inhibitor cocktail (MilliporeSigma). Cell lysate was incubated on ice for 30 minutes, vortexed every 10 minutes, and then centrifuged at 12,000 rpm for 20 minutes at 4°C. The supernatant was collected and measured for protein concentration using Pierce BCA Protein Assay Kit (Thermo Scientific). For Western blot, a total of 20 µg whole cell lysate was resolved in a 10 % precast Mini-PROTEAN TGX Stain-Free gel (Bio-Rad). At the end of the run, the gel was exposed to UV light for 2.5 minutes in Bio-Rad ChemiDoc XRS+ System to activate the trihalo compound that is bound to tryptophan residues. Proteins were then transferred from gel into Immobilon-P PVDF membrane (MilliporeSigma). A fluorescent stain-free blot image was captured using Bio-Rad ChemiDoc XRS+ System as the total protein signal for later normalization of protein loading. The membrane was blocked using Chemi Blot Blocking Buffer (Azure Biosystems) for 1 hour at room temperature with gentle shaking. The membrane was subsequently cut into two parts. The upper portion was incubated with a mouse monoclonal antibody against YY1 (Santa Cruz Biotechnologies #sc-7341X) at 1:10,000 dilution and the lower portion was incubated with a mouse monoclonal antibody against histone H3 protein (Santa Cruz Biotechnologies #sc-517576) at 1:5,000 dilution. After overnight incubation at 4 °C with primary antibodies, the membrane was incubated with an HRP conjugated goat anti-mouse IgG secondary antibody (Bio-Rad # 1706516) for 1 hour at room temperature. Chemiluminescent signal was generated by adding ECL Select Western blot detecting reagent (Cytiva Life Sciences) and imaged with Bio-Rad ChemiDoc XRS+ System. Bio-Rad Image Labs v4.0 was used to quantify the YY1 protein signals and normalized to either total protein signals (e.g., from the fluorescent Stain-Free blot image) or the housekeeping H3 protein signals.

### Electrophoretic mobility shift assay (EMSA)

Details about all DNA fragments used in EMSA are provided as a supplemental table (Table S2). EMSA was performed using LightShift Chemiluminescent EMSA Kit (Thermo Scientific). Nuclear proteins were extracted using NE-PER Nuclear and Cytoplasmic Extraction Reagents (Thermo Scientific). In brief, F9 or NIH/3T3 cells at 90% confluence were trypsinized, centrifuged for 5 minutes at 500 g, and washed with 1x PBS. The pellet was resuspended in ice-cold cytoplasmic extraction reagent with protease inhibitor and centrifuged. The supernatant containing cytoplasmic proteins was removed. The pellet was resuspended in ice-cold nuclear extraction reagent with protease inhibitor, incubated on ice for a total of 40 minutes while vortexing for 15 seconds every 10 minutes, and centrifuged for 10 minutes at 16,000 g at 4°C. The supernatant containing nuclear proteins was measured for protein concentration using Pierce BCA Protein Assay Kit (Thermo Scientific) and stored at -80°C until use. To generate biotin-labeled double-stranded DNA probe, a 5’ biotinylated 30-mer oligo and an unlabeled reverse complement oligo were annealed at a 1 µM concentration in 1 x TE buffer in the presence of 50 mM NaCl. Unlabeled competitor DNA fragments were generated in the same fashion but at a concentration of 2 µM. Binding reactions were set up in a 20 µL volume containing 1x binding buffer, 2.5% glycerol, 5 mM MgCl2, 50 ng/mL poly dI dC, 0.05% NP-40, 20 fmol of annealed probe in the presence or absence of unlabeled competitor, and 4.5 µg of nuclear protein extract from either NIH/3T3 or F9 cells. Binding reactions were incubated for 20 minutes at room temperature without antibodies. After the binding reaction and when indicated, 2.5 µg of YY1 specific antibody (Santa Cruz Biotechnologies #sc-7341X) or mouse IgG isotype control (Thermo Fisher #02-6502) were added and incubated at room temperature for 20 more minutes. A 5% polyacrylamide gel was pre-run in 0.5x TBE for 30 minutes. Each reaction was mixed with 5 µL of 5x loading dye, loaded, and run on for 1.5 hour at 100 volts. Samples were then transferred to a nylon membrane (pre-soaked in cold 0.5x TBE for 10 minutes) at 380 mA for 30 minutes at 10°C. DNA was crosslinked to the membrane using Stratagene UV crosslinker 1800 instrument at 120mJ/cm2 (using the auto crosslink function). The membrane was blocked for 15 minutes using blocking buffer and incubated in conjugate/blocking buffer for 15 minutes. The membrane was washed four times for 5 minutes each, and equilibrated for 5 minutes in substrate equilibration buffer, and incubated in substrate working solution for 5 minutes. The chemiluminescence was captured by Bio-Rad ChemiDoc XRS+ System.

### Enrichment analysis of YY1 ChIP-seq data across TE subfamilies using T3E

Raw YY1 ChIP-seq data (39) were downloaded from NCBI Gene Expression Omnibus (GEO). The following ChIP-seq datasets were used for enrichment analysis: three replicates of untreated (i.e., treated with DMSO vehicle) wild-type mESC J1 strain immunoprecipitated by YY1 antibody (Santa Cruz, sc-1703) (GEO sample ID: GSM3772791, GSM3772795 and GSM3772799) and the corresponding input sample (GEO sample ID: GSM3772809). All four libraries consisted of 75 bp paired-end reads from Illumina HiSeq4000 platform. Reads were preprocessed as previously described for T3E (40). Low-quality bases (Phred score < 20) and adapter sequences were trimmed from the 3’ end of the ChIP-seq reads. Sequencing reads were aligned to the mouse reference genome (GRCm38/mm10) using BWA-MEM v0.7.17 (41) with parameter “-a”, returning all mappings (both unimappers and multimappers). Duplicate reads, un-mapped reads, and alignments to the mitochondrial chromosome and non-chromosomal scaffolds were removed. Resulting alignment files of three replicates and one input control were fed into T3E (40) for enrichment analysis. T3E repository was cloned from GitHub (https://github.com/michelleapaz/T3E) and installed on the institutional high-performance computing Linux cluster. Repeat annotations were derived from mm10 repeat library db20140131, available from the RepeatMasker website (http://www.repeatmasker.org), and filtered to retain only individual instances of TEs (i.e., LINE, SINE, LTR and DNA transposons). T3E calculates enrichment for TE subfamilies, not for individual TE copies, returning a fold-change (FC) and an empirical P value. For a given TE subfamily, P value < 0.01 is considered enrichment for the protein of interest. Out of 1159 TE subfamilies, 118 are enriched for YY1 binding (p < 0.01) across all three ChIP-seq samples. FC_mean is the mean fold change among three replicates (the average is taken after converting each log2FC into FC) (Table S3).

## Results

### Mutating the predicted YY1-binding site abolishes Tf_I promoter activity in reporter assays

In the 5’UTR of mouse L1 Tf subfamily, a YY1-binding motif was predicted at nt 77-88 (GTC<u>GCCAT</u>CTTG) in the monomer consensus (31) (Fig.1A; motif 1). It contains the five core nucleotides (GCCAT) that are highly conserved among mouse and human YY1-binding sites (42) (Fig.1B). To evaluate the function of this putative YY1-binding site, we employed a single-vector dual-luciferase reporter assay as previously reported in mouse F9 embryonal carcinoma cells (34). In this reporter assay, the firefly luciferase (Fluc) is controlled by an L1 promoter variant. The Renilla luciferase (Rluc) is driven by herpes simplex virus thymidine kinase (HSV-TK) promoter and used to normalize transfection efficiency. To quantify the activity of an L1 promoter variant, Fluc/Rluc ratios are first calculated for each of the four replicate wells and then averaged. An average Fluc/Rluc ratio is similarly calculated for a negative control plasmid pLK037, which lacks a promoter sequence upstream of the Fluc coding sequence (34), representing the assay background. The average Fluc/Rluc ratio of each promoter construct is subsequently normalized to that of pLK037 (i.e., setting the average Fluc/Rluc ratio of pLK037 to 1), giving rise to “normalized promoter activity” (Fig.1C). Each experiment also includes a positive control plasmid pCH117, which contains a highly active human L1 promoter upstream of the Fluc coding sequence (34). The normalized promoter activity for pCH117 represents the assay dynamic range and is stated in the figure legend for each experiment.

**Figure 1.**
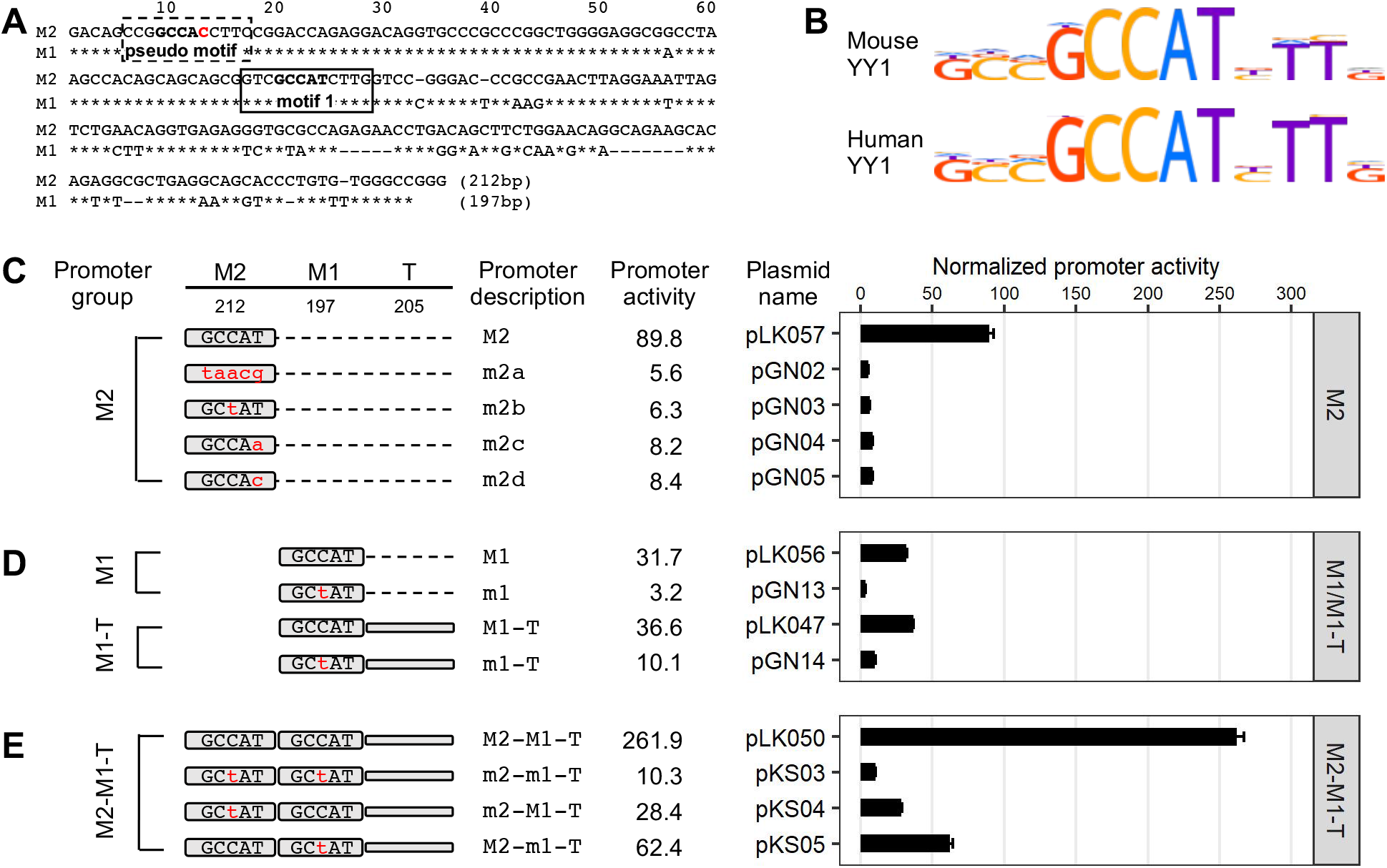
Effect of YY1 motif mutants on Tf_I promoter activity in F9 cells. **(A)** Alignment of Tf_I monomer 2 (M2) and monomer 1 (M1) consensus sequences. In the M1 sequence, nucleotide positions identical to M2 are marked by asterisks. Sequence gaps are represented by dashes. A previously predicted YY1 binding motif is located between nt 77-88 (solid box, termed “motif 1”). Toward the 5’ end of the monomers is another stretch of nucleotides highly similar to the consensus YY1 binding motif (dashed box, termed “pseudo motif”). **(B)** Mouse and human YY1 binding motif models from the HOCOMOCO database. **(C)** Normalized promoter activity of M2 constructs. Mutation to consensus YY1 binding motif 1 (GCCAT) is indicated by lowercases in red. **(D)** Normalized promoter activity of M1 and M1-Tether (M1-T) constructs. **(E)** Normalized promoter activity of M2-M1-T constructs. Mutation to one monomer at a time showed activity from the other monomer and tether. For panels C-E, sequence organization of the promoters is illustrated on the left side. The length of M2, M1, and tether for each promoter is annotated (in base pairs). The dashed line represents domain(s) that were removed in reference to the two-monomer 5’UTR sequence (M2-M1-T). The x-axis indicates the normalized promoter activity, which is also listed under column “promoter activity” for each promoter variant. The positive control construct, pCH117, had a normalized promoter activity of 355.0. Error bars represent standard errors of the mean (n = 4).

In the first experiment, we compared the consensus monomer 2 (M2) of Tf_I subfamily to four variants containing mutated YY1-binding sites (Fig.1C). The four mutant variants had either one or all five core nucleotides altered as compared with the core consensus YY1-binding sequence GCCAT. Three of the four variants had been previously shown to behave as a loss-of-function mutation in other promoter contexts: taacg (variant m2a; lowercase indicates substitution) (43), GCtAT (variant m2b) (44), GCCAa (variant m2c) (45). The fourth variant, GCCAc (m2d), was similar to variant m2c but had a T to C transition at the fifth nucleotide position instead. As expected, Tf_I M2 showed a normalized promoter activity of 89.8 (Fig. 1C). In contrast, the four mutant promoter variants uniformly displayed minimal promoter activity, ranging from 5.6 to 8.4 and corresponding to 10.7 to 16.0-fold reduction as compared to the wild-type M2. The mutant promoters’ significant loss of activity suggests that the putative YY1-binding sequence is essential for transcriptional activation of Tf_I M2 when tested in isolation (i.e., when not linked to downstream 5’UTR sequence).

In the second experiment, we tested the function of the putative YY1-binding site in the context of monomer 1 (M1) alone by introducing a single, centrally located nucleotide substitution as in m2b (Fig.1D). The mutant monomer 1 (m1) showed 10-fold less activity than the wild-type M1. Similarly, in the context of monomer 1 followed by the tether sequence (M1-T), the mutant version (m1-T) had 3.6-fold reduced activity than the wild-type (Fig.1D). The higher residual activity seen in m1-T reflects the inherent contribution from the tether sequence (34).

In the third experiment, we conducted mutational analysis in the context of two-monomer Tf_I 5’UTR (M2-M1-T) (Fig.1E). As expected, the consensus M2-M1-T possessed significantly higher activity than M2 alone, M1 alone, or M1-T (Fig.1C-E), reproducing the synergistic interaction among M2, M1 and T seen earlier (34). When the putative YY1 motif was mutated in both monomers (m2-m1-T), the promoter activity was reduced to 10.3, equivalent to that of m1-T. This observation suggests that the synergy among M2, M1 and T stems predominantly from the presence of the putative YY1 motifs in M2 and M1. Indeed, a singular mutation in M2 (m2-M1-T) reduced the activity to that of M1-T (28.4 versus 36.6, respectively). A singular mutation in M1 (M2-m1-T; Fig.1E) reduced its activity to a level that was similar to M2 alone (Fig.1C) (62.4 versus 89.8, respectively).

To exclude cell-specific artifacts, we repeated the experiments in the NIH/3T3 mouse embryonic fibroblast cell line and observed similar results (Fig. S1). Briefly, the four mutant M2 promoter variants uniformly displayed minimal activity, corresponding to 16-to 22-fold reduction relative to the wild-type (Fig.S1A). Variant m1 and m1-T showed 11-fold and 2.4-fold reduction in activity relative to their wild-type counterparts (Fig.S1B). When either or both YY1-binding motifs were mutated in the context of M2-M1-T promoter constructs a synergistic interaction between M2 and M1 was also reproduced (Fig.S1C). Taken together, these reporter assays suggest that the previously predicted YY1-binding motif in the consensus Tf_I monomer sequences is not only critical for each monomer’s own promoter activity but also responsible for the synergistic interaction between monomers.

### The putative YY-binding motif in Tf_I 5’UTR interacts with YY1 protein

To determine whether the putative YY1-binding motif in Tf_I 5’UTR interacts with YY1 protein, we utilized the electrophoretic mobility shift assay (EMSA). As a probe for motif 1, we used a biotin-labeled 30 bp double-stranded DNA fragment from Tf_1 M2, with motif 1 centrally located (Fig.2A; WT fragment). Incubation with nuclear protein extract from F9 cells resulted in a shift in its migration (Fig.2B, lane 2). A supershift was observed with the addition of YY1-specific antibody (Fig.2B, lane 3) but not with the addition of non-specific mouse IgG (Fig.2B, lane 4), suggesting the interaction is mediated by YY1 protein. The shift was diminished by increasing amount of unlabeled DNA of the same sequence (Fig.2B, lanes 5-7) but not by three unlabeled DNA fragments (Mut1, Mut2, and Mut3 fragments shown in Fig.2A in which the core nucleotides were variably mutated as in the reporter assays) (Fig.2B, lanes 8-10). Similar results were obtained when nuclear protein extract from NIH/3T3 cells was used (Fig.S2B). The inability of these mutant DNA fragments to compete for YY1 binding highlights the presence of sequence-specific interaction of the predicted binding motif with YY1 protein.

**Figure 2.**
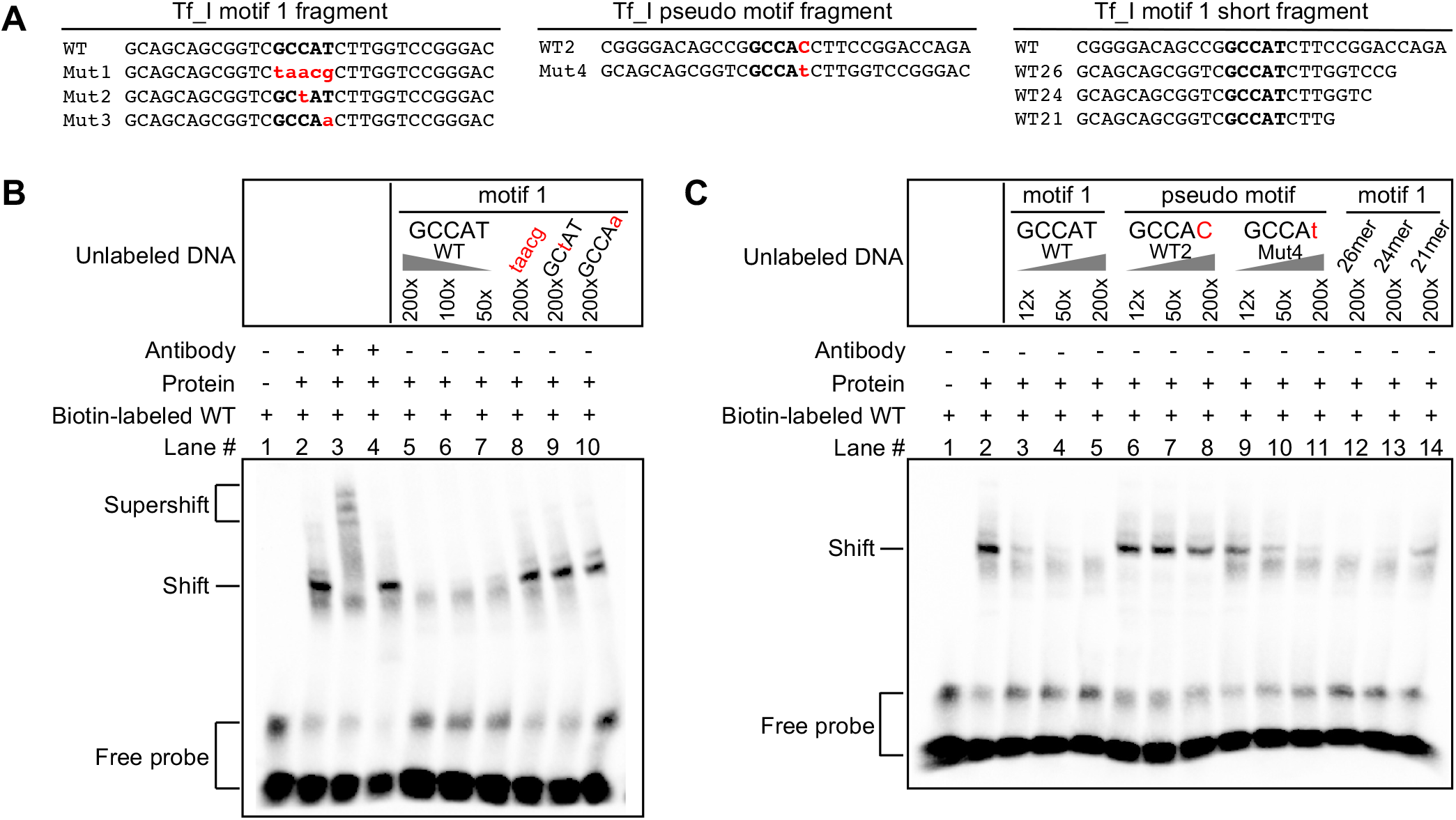
Interaction of YY1 protein with motif 1 but not with a pseudo motif in Tf_I monomers. (**A**) Wild-type and mutant DNA fragments used in electrophoretic mobility shift assays (EMSA). Each fragment was formed by annealing a sense stranded oligo (shown) with the corresponding antisense oligo (not shown). Mutations in the core binding motif are indicated by lowercases in red. The wild-type pseudo motif (WT2) differs from motif 1 (WT) at the fifth position of the core binding sequence. (**B**) EMSA with Tf_I motif 1 fragments. The presence or absence of a biotin-labeled WT probe, antibody, and nuclear protein extract from F9 cells is indicated by “+” or “-” symbols. Lane 3 had YY1-specific antibody and lane 4 had mouse IgG as a control. Lanes 5-10 had unlabeled DNA fragments as competitors in molar excess as indicated. (**C**) EMSA with Tf_I pseudo motif fragments and shortened motif 1 fragments. The presence or absence of a biotin-labeled WT probe, antibody, and nuclear protein extract from F9 cells is indicated by “+” or “-” symbols. Lanes 3-14 had unlabeled DNA fragments as competitors in molar excess as indicated.

In addition to motif 1 that was predicted by earlier studies, we noted the presence of another string of nucleotides at nt 6-17 (CCG<u>GCCAC</u>CTTC) that closely resembles the consensus YY1-binding site. This motif is conserved at four out of the five core nucleotides (GCCAC instead of GCCAT). As tested below, it is unable to interact with YY1 protein, so we named it “pseudo motif” (Fig.1A). To determine whether the pseudo motif interacts with YY1 protein we synthesized two unlabeled 30 bp DNA fragments for EMSA: a wild-type DNA fragment based on Tf_I M2, with the pseudo motif centrally located (Fig.2A, WT2 GCCAC), and a mutant version with a single nucleotide substitution that restores it to the consensus YY1-binding site at the fifth core nucleotide position (Fig.2A, Mut4 GCCAt). For this EMSA trial, because we were uncertain whether a biotin-labeled pseudo motif fragment would bind to YY1 protein and yield a shift signal, we decided to keep the biotin-labeled motif 1 fragment as the probe. In this design, binding of a DNA fragment to YY1 was evaluated by its potency as an unlabeled competitor. As a control (Fig.2C, lanes 3-5), unlabeled motif 1 fragment diminished the shift in a quantitative manner: a significant reduction even at a 12-fold excess of unlabeled competitors and a 50-fold excess nearly eliminated probe binding. In contrast, the wild-type pseudo motif fragment could hardly compete with probe binding (Fig.2C, lanes 6-8): some level of competition could only be discerned at 200-fold excess. Interestingly, unlabeled Mut4 fragment was able to reduce the shift in a dose-dependent manner although not as potent as the unlabeled motif 1 fragment (Fig.2C, lanes 9-11). Similar results were obtained when nuclear protein extract from NIH/3T3 cells was used (Fig.S2C). These results suggest that the pseudo motif is unable to interact with YY1 protein due to its single nucleotide deviation from the consensus 5-nt core motif.

So far, we have used 30 bp DNA fragments as probes or competitors in EMSA trials. To evaluate the effect of sequence flanking the binding motif on YY1 protein binding, we progressively shortened the DNA fragments from the 3’ end, creating 26, 24 and 21 bp fragments (Fig.2A; WT26, WT24 and WT21). When used as unlabeled competitors, the 26 bp and 24 bp fragments were similarly effective as compared to the 30 bp WT fragment (Fig.2C, lanes 12-13). On the other hand, the 21 bp fragment was much less effective at 200-fold excess and only achieved the level of inhibition equivalent to the 30 bp WT fragment at 12-fold excess (Fig.2C, compare lands 3 and 14). These results suggest that the extra nucleotides flanking the YY1-binding motif potentiate its interaction with YY1 protein in vitro. Similar results were observed when the experiment was repeated with nuclear protein extract from NIH/3T3 cells (Fig.S2B).

### Knockdown of YY1 protein by siRNA reduces Tf_I promoter activity

So far, we have provided evidence that Tf_I 5’UTR loses its promoter activity when motif 1 is mutated (Fig.1) and motif 1 mediates the interaction between YY1 protein and Tf_I monomers (Fig.2). If YY1 protein binding to motif 1 is responsible for Tf_I promoter activity, a decrease in YY1 protein abundance should lead to reduced transcriptional activation. To knock down YY1 protein, we first evaluated four siRNAs against mouse Yy1 RNA (Yy1_1, Yy1_5, Yy1_6, and Yy1_7) in NIH/3T3 cells by Western blot (Fig.3A). Note the protein has an apparent molecular weight of 68 kD in despite having a calculated molecular weight of 44.7 kD (17). As compared to a control non-specific siRNA, the highest knockdown efficiency was achieved by Yy1_1 (89%), followed by Yy1_7 (84%), Yy1_5 (78%) and Yy1_6 (33%). A knockdown efficiency of 88% was achieved when all four siRNAs were pooled together. Subsequently, we selected siRNA Yy1_1 and Yy1_7 for three Tf_I promoter assays (M2, M1, or M2-M1-T) in NIH/3T3 cells (Fig.3B). For each promoter group, its normalized promoter activity was largely unaffected by the negative control non-specific siRNA (Fig.3B; compare Allstars with no siRNA). In reference to Allstars, Yy1_1 siRNA treated cells showed 34.3%, 30.1%, and 36.7% of the activity for M2, M1, and M2-M1-T, respectively. In comparison, Yy1_7 treated cells showed 44.7%, 46.7%, and 54.0% of the activity for M2, M1, and M2-M1-T, respectively. When repeated in F9 cells, Yy1_1 siRNA treatment reduced M2, M1, and M2-M1-T activities to 46.3%, 60.7%, and 43.5%, respectively; Yy1_7 siRNA decreased their activities to 56.1%, 71.4%, and 51.7%, respectively (Fig.S3). These results confirm that YY1 functions as a transcriptional activator for Tf_I monomers in a cell-based reporter assay.

**Figure 3.**
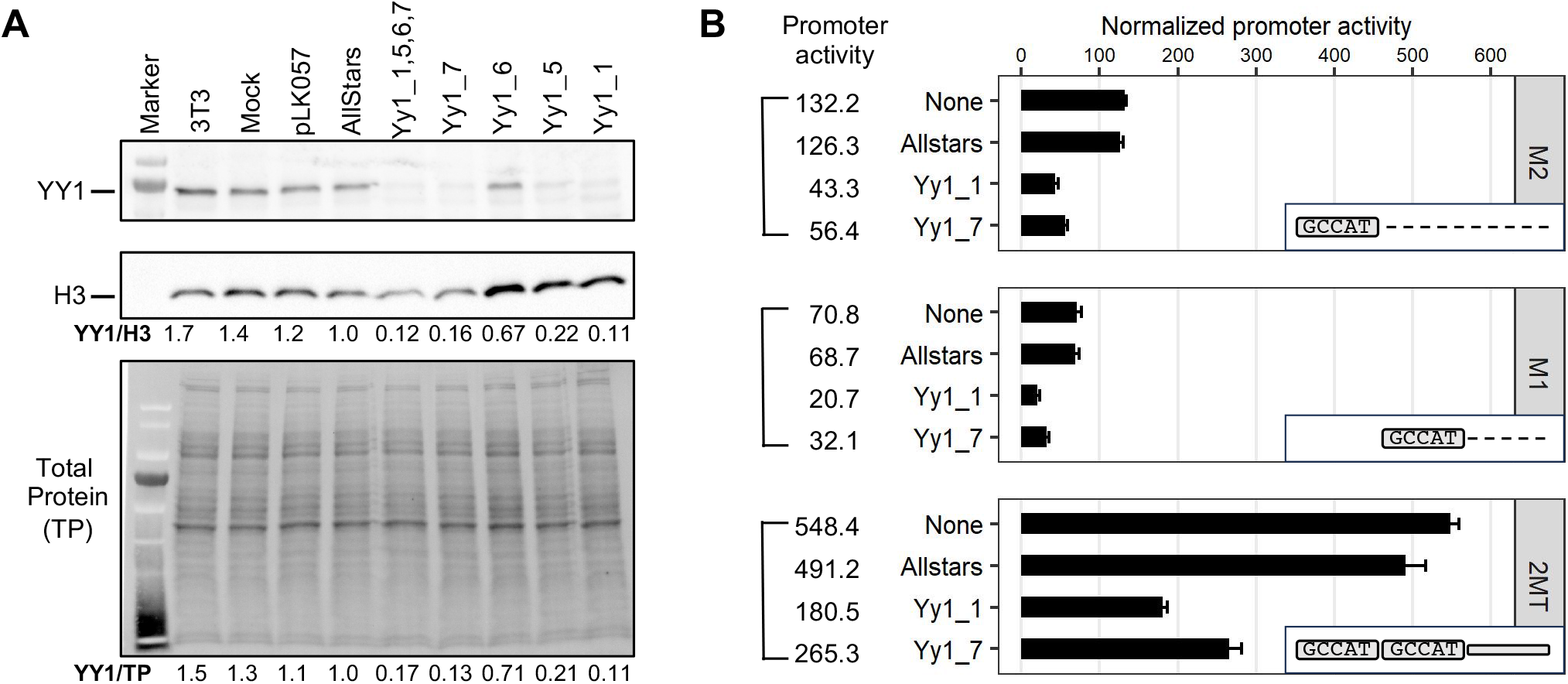
Knockdown of YY1 protein and its impact on the Tf_I promoter activity. **(A)** Western blot analysis of YY1 protein knockdown. NIH/3T3 cells were transfected with four YY1-specific siRNAs individually or as a pool. After 72 hours whole cell lysates were probed for YY1 protein (top panel). Yy1_1 and Yy1_7 showed efficient knockdown compared to the control siRNA (Allstars). The YY1 protein signal was normalized to either histone H3 (middle panel) or fluorescent stain-free total protein signal (lower panel). (**B)** Normalized promoter activity for three Tf_I promoter constructs (M2, M1, and M2-M1-T) under siRNA knockdown. For each promoter variant, cells were cotransfected with the promoter construct and with or without a siRNA (y axis; none = no siRNA). In reference to cells treated with Allstars, Yy1_1 siRNA treated cells showed 34.3%, 30.1%, and 36.7% of the activity for M2, M1, and M2-M1-T, respectively. In comparison, Yy1_7 treated cells showed 44.7%, 46.7%, and 54.0% of the activity for M2, M1, and M2-M1-T (marked as 2MT), respectively. The positive control construct, pCH117, had a normalized promoter activity of 1222.8. Error bars represent standard errors of the mean (n = 4). The inset illustrates the promoter constructs used.

### Gf_I monomers lack functional YY1-binding motif

The Gf subfamily of mouse L1s was first reported in 2001 (36). The consensus Gf monomer shares sequence homology with Tf monomers. At the position corresponding to Tf_I motif 1, Gf monomer contains sequence GGA<u>GCCTT</u>CTTG, which deviates from the 5-nt consensus YY1-binding motif by one nucleotide (GCCTT instead of GCCAT) (Fig.4A). To determine whether this motif is important for Gf_I promoter activity, we created and tested two mutant variants for both Gf_I M2 and M1 in reporter assays in F9 cells (Fig.4B-C). The first mutant variant had all five core nucleotides altered (m2a and m1a, “taacg”). The same alteration completely abolished the promoter activity of Tf_I monomers in earlier experiments. However, minimal change in promoter activity was observed in the context of both Gf_I monomers (Fig.4B-C; compare m2a with M2 and compare m1a with M1), suggesting the original GCCTT motif in Gf_I monomers is not involved in transcriptional activation or repression. In the second mutant variant, we introduced a nucleotide substitution that effectively converted the original motif into a consensus YY1 core motif (m2b and m1b, “GCCaT”). Interestingly, this single nucleotide change elevated the Gf_I M2 promoter activity by 29.8-fold (Fig.4B, compare m2b with M2), even more active than the Tf_I M2 fragment (see Fig.1C). The same nucleotide change enhanced the M1 activity by 6.6-fold (Fig.4C, compare m1b with M1). Lastly, we introduced this nucleotide change to both monomers in the context of M2-M1-T and observed a significant 3.5-fold boost to the promoter activity (Fig.4D). Similar results were obtained from reporter assays conducted in NIH/3T3 cells (Fig.S4). Variant m2b, m1b, and m2-m1-T showed 36.1-, 7.0-, and 6.1-fold higher activities, respectively, than their wild-type counterparts. These data support the conclusion that the single nucleotide divergence in Gf_I monomers negatively affects its transcriptional output.

**Figure 4.**
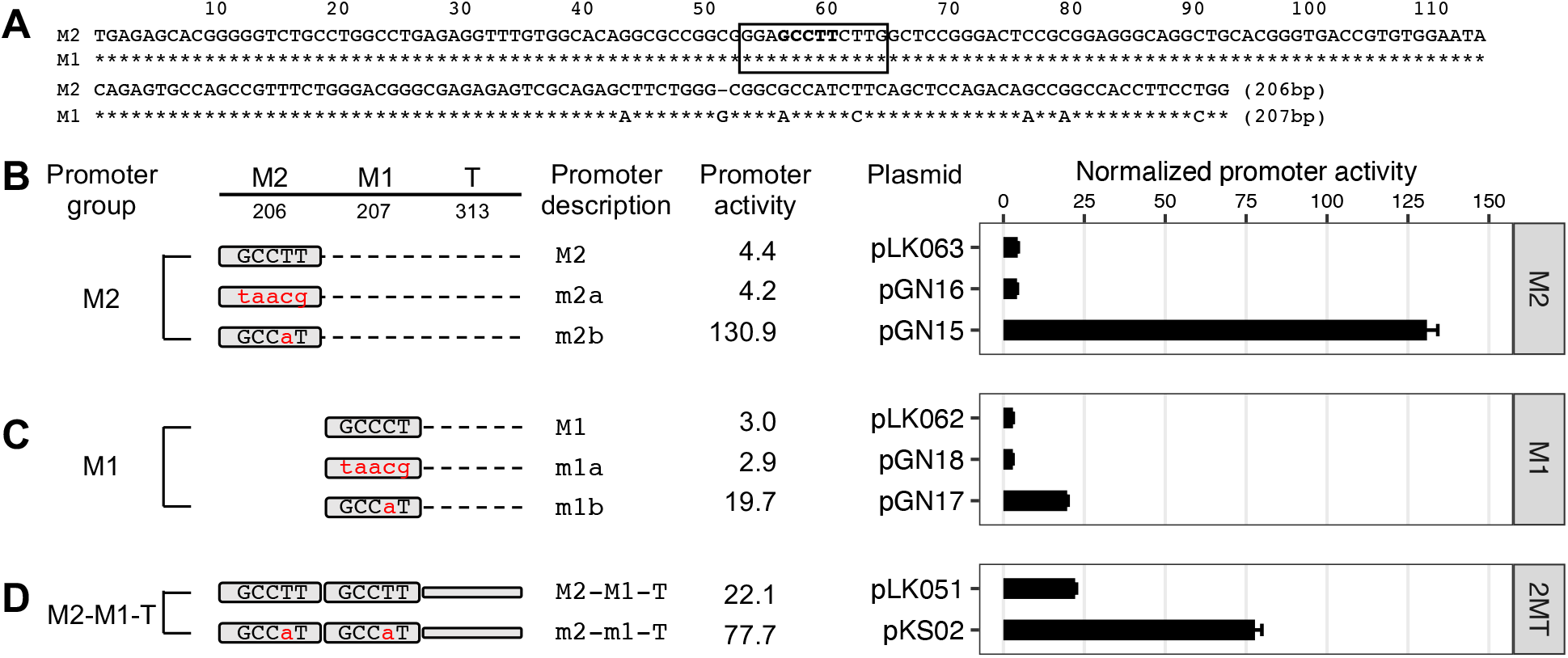
Promoter activities of YY1 motif variants of the Gf_I subfamily in F9 cells. **(A)** Alignment of Gf_I monomer 2 (M2) and monomer 1 (M1) consensus sequences. In the M1 sequence, nucleotide positions identical to M2 are marked by asterisks. Sequence gaps are represented by dashes. A previously predicted YY1 binding motif is located between nt 53-64 (solid box, termed “Gf_I motif”). Promoter activity is assessed using Dual-luciferase reporter assay. **(B)** Normalized promoter activity of monomer2 (M2) constructs. Mutation to Gf_I motif (GCCCT) is indicated by lowercases in red. The m2a showed minimal change in promoter activity. However, changing to the consensus YY1 motif (m2b) elevated the M2 promoter activity by 29.8-fold. **(C)** Normalized promoter activity of monomer 1 (M1) constructs. The mutant monomer 1 (m1a) showed minimal change in promoter activity. Changing to the consensus (m1b) showed 6.6 times higher signal compared to M1. **(D)** Normalized promoter activity of monomer2-monomer1-Tether (M2-M1-T; or 2MT) constructs. A 3.5-fold higher activity was observed upon changing both Gf_I motifs to the consensus sequence. The positive control construct, pCH117, had a normalized promoter activity of 493.0. Error bars represent standard errors of the mean (n = 4).

To explore whether these functional changes are mediated by YY1 transcription factor, we tested the ability of the wild-type and corrected Gf_I motif to interact with YY1 protein using EMSA. We synthesized two 30 bp DNA fragments for EMSA: a wild-type DNA fragment based on Gf_I M2, with the putative motif centrally located (Fig.5A, WT GCCTT), and a mutant version with a single nucleotide substitution at the fourth core nucleotide position that restores it to the consensus YY1-binding site (Fig.5A, Mut GCCaT). Again, as we were uncertain whether a biotin-labeled WT motif fragment would bind to YY1-containing nuclear extract from F9 cells and yield a shift signal, we used the biotin-labeled Mut fragment as the probe. Indeed, nuclear protein extract bound to the GCCaT probe, causing a shift (Fig.5B, lane 2). A supershift was observed with the addition of YY1 antibody (Fig.5B, lane 3) but not with the addition of non-specific mouse IgG (Fig.5B, lane 4), confirming the involvement of YY1 protein. The shift was diminished by an excess of unlabeled GCCaT DNA (Fig.5B, lanes 5-7) but not by the WT DNA fragment (Fig.5B, lanes 8-10). The inability of WT DNA fragments to compete for YY1 binding indicates that the single nucleotide difference prevents its interaction with YY1 protein. Similar results were obtained when nuclear protein extract from NIH/3T3 cells was used (Fig.S5). Together, these results suggest that the homologous region in Gf_I monomers is unable to interact with YY1 protein due to its single nucleotide deviation from the consensus 5-nt core motif.

**Figure 5.**
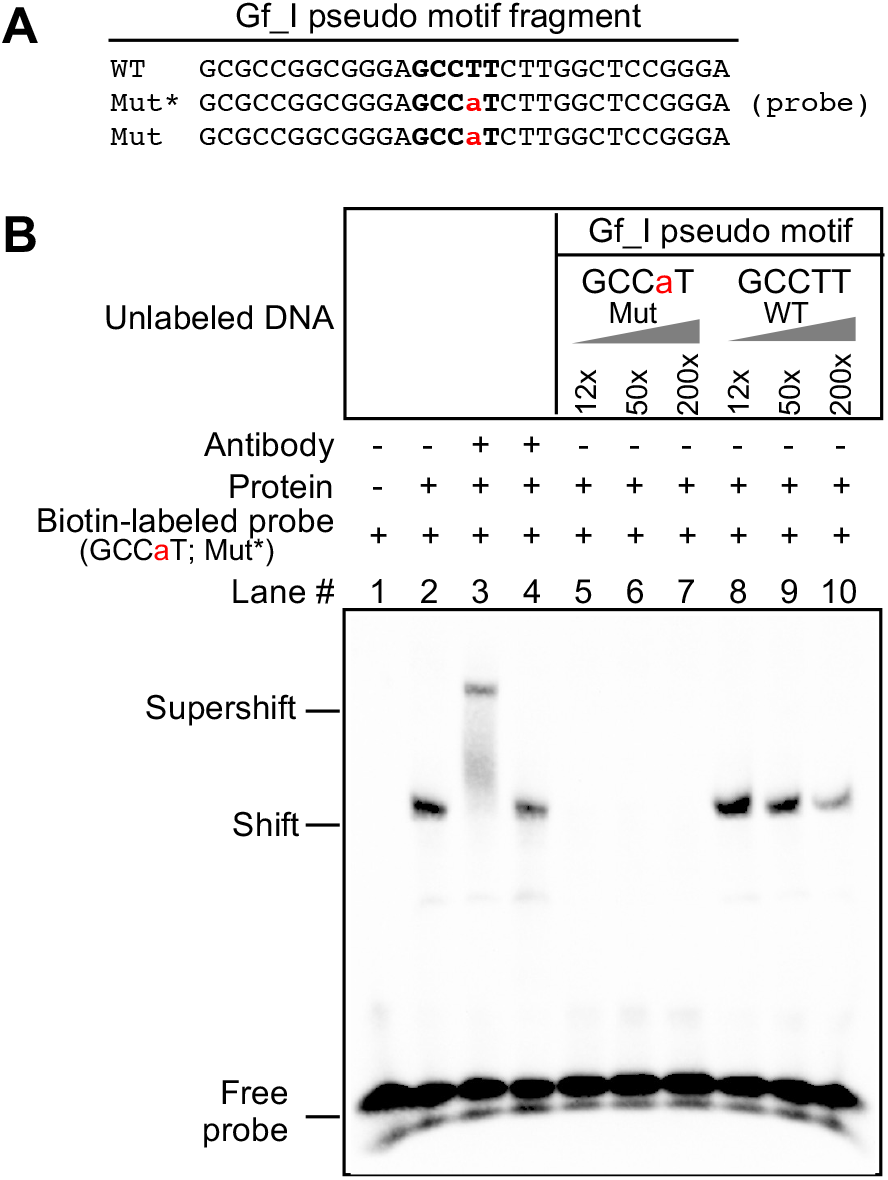
Lack of interaction of YY1-containing nuclear protein extract with a putative YY1 binding motif in Gf_I monomers. **(A)** Wild-type and mutant DNA fragments used in electrophoretic mobility shift assays (EMSA). Each fragment was formed by annealing a sense stranded oligo (shown) with the corresponding antisense oligo (not shown). Mutation in the core binding motif is indicated by a lowercase letter in red. In the mutant variant (Mut), the wild-type pseudo motif (WT) was mutated at the fourth position to restore the consensus core binding sequence. **(B)** EMSA with Gf_I pseudo motif fragments. The presence or absence of a biotin-labeled Mut probe (GCCaT), antibody, and nuclear protein extract from F9 cells is indicated by “+” or “-” symbols. Lane 3 had YY1-specific antibody and lane 4 had mouse IgG as a control. Lanes 5-10 had unlabeled DNA fragments as competitors in molar excess as indicated.

### Tf_I and Tf_II subfamilies are enriched for YY1 binding in mouse embryonic stem cells

It has been shown previously that YY1 occupancy is increased in mouse embryonic stem cells (mESCs) upon loss of DNA methylation and histone deacetylase activity (39). Of interest, 652 out of 1067 (61%) YY1-binding sites gained overlap with LINEs, predominantly with Tf subfamily (537 out of 652) followed by Gf subfamily (54 out of 652) (39). However, the steady-state YY1 distribution across TE subfamilies in wild-type mESCs has not been reported. To compare YY1 occupancy among TE subfamilies we utilized Transposable Element Enrichment Estimator (T3E), a recently published ChIP-seq analysis pipeline specifically designed to profile protein binding across TE families/subfamilies (40). T3E computes the number of read mappings for a TE subfamily by counting both unique- and multiple-mapped reads. Among 1159 TE subfamilies encompassing LINE, SINE, LTR retrotransposons and DNA transposons, 10% (118/1159) were found enriched for YY1 binding (p < 0.01) across all three ChIP-seq samples (Fig.6A; Table S3). The degree of enrichment is reflected by fold-change, a ratio between read mappings in the ChIP-seq sample and the average of simulations based on the input library. Only three TE subfamilies displayed two-fold or higher fold-change. RLTR43A, a member of LTR/ERVK family with only three genomic loci, has the highest average fold-change (7.5) among the three ChIP-seq samples. The second- and third-ranked subfamilies are Tf_I and Tf_II (2.2 and 2.0, respectively) (Fig.6B). In total, 11 LINE subfamilies showed enrichment for YY1 (Fig.6C-D). However, none of the other evolutionarily young mouse L1 subfamilies were enriched for YY1 binding, such as Tf_III, Gf_I, A_I, A_II, and N_I (Figure 6B).

**Figure 6.**
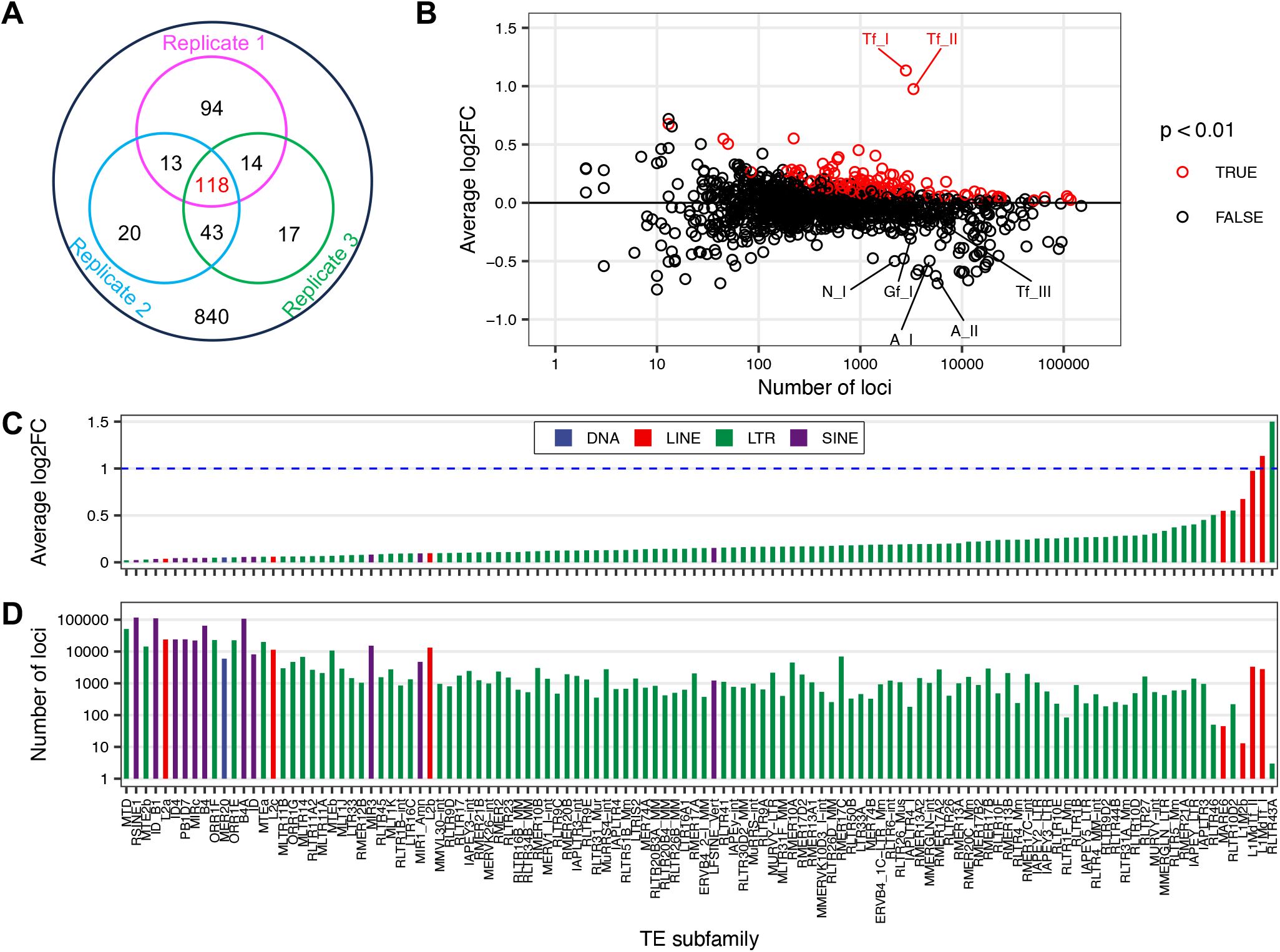
Enrichment of YY1 binding among TE subfamilies in mouse embryonic stem cells. (**A**) Distribution of TE subfamilies enriched for YY1 binding in one, two or all three replicate ChIP-seq samples. Each circle corresponds to the number of TE subfamilies enriched for YY1 binding in one of the three sample (p < 0.01). (**B**) Enrichment analysis of YY1 binding across 1159 TE subfamilies. The log2 fold-change (log2FC, y axis) for each subfamily is plotted against its number of loci in the mouse genome. TE subfamilies with p value < 0.01 (red circles) are considered enriched for YY1 binding. Seven evolutionarily young L1 subfamilies are labelled. Note the top ranked subfamily RLTR43A (with log2FC of 2.9 and 3 genomic loci) is not shown. (**C**) Average log2FC among three replicate ChIP-seq samples across 118 TE subfamilies enriched for YY1 binding (p < 0.01). Subfamilies are color coded by TE classes and arranged by log2FC in ascending order on the x-axis. Panel C shares x-axis with panel D. Note the log2FC value for the top ranked subfamily RLTR43A is truncated at 1.5 on the graph. The dashed blue line marks 2-fold change (i.e., log2FC = 1). (**D**) The number of genomic loci for TE subfamilies shown in panel C.

In addition to 7 LINE subfamilies, 99 LTR subfamilies, 11 SINE subfamilies, and one subfamily of DNA transposon showed varied enrichment of YY1 binding in mESCs (Fig.6C-D). Three families (ERV1, ERV2/ERVK, and ERV3/ERVL) make up mouse LTR class of TEs (46), and they contributed to 22, 60 and 17 subfamilies enriched for YY1 occupancy, respectively but with modest fold changes. Intracisternal A-type particle (IAP) subfamilies belong to family ERV2/ERVK. YY1-binding site has been previously found in the promoter region of IAP (47,48). Consistent with these studies, ten IAP subfamilies showed YY1 enrichment with fold changes ranging from 1.1 to 1.4 (Fig.6C-D). Enrichment of YY1 motif in ERV3/MERVL has also been reported (48,49).

## Discussion

Unlike human L1, the role of YY1 in mouse L1 transcription had not been experimentally tested before. In this study, we examined YY1’s function in the transcription of two mouse L1 subfamilies, known as Tf and Gf when they were initially discovered (30,36). We provided multiple lines of evidence that support an activating role of YY1 for Tf subfamilies: (i) In luciferase-based reporter assays, mutating the conserved nucleotides of the putative YY1-binding site diminished the promoter activity in four different promoter constructs (M2 alone, M1 alone, M1-T, and M2-M1-T) (Fig.1C-E and Fig.S1). (ii) In EMSA, 30 bp DNA fragments containing the putative YY1-binding motif showed sequence-specific interaction with YY1-containing nuclear protein extract (Fig.2B and Fig.S2B). (iii) siRNA knockdown of YY1 protein led to reduced promoter activities in reporter assays (Fig.3B and Fig.S3). In parallel, we provided experimental evidence that excluded a role of YY1 for Gf subfamily: (i) In reporter assays, mutating all five core nucleotides in the putative YY1-binding motif had no effect on activity of M2 or M1. In contrast, a single nucleotide substitution restoring to the consensus led to multi-folds of increase in promoter activity for three promoter constructs (M2, M1, and M2-M1-T) (Fig.4 and Fig.S4). (ii) In EMSA, while DNA fragments containing the single nucleotide substitution interacted with YY1-containing nuclear protein extract, DNA fragments containing the putative YY1-binding motif failed when used as a competitor (Fig.5 and Fig.S5). These data indicate that the lack of YY1 binding is the result of a single nucleotide deviation in Gf promoter (GCCTT) from the consensus YY1-binding motif (GCCAT). Additionally, we excluded the presence of a second YY1-interacting motif in Tf monomers. This motif (the pseudo motif) bears a GCCAC core sequence, also a single nucleotide deviation from the YY1 consensus (GCCAT). We showed that (i) In EMSA, DNA fragments containing the wild-type GCCAC motif could not compete for YY1 binding. DNA fragments containing a single nucleotide substitution restoring it to the consensus had enhanced interaction with YY1-containing nuclear protein extract although not as efficient as the motif 1 containing DNA (Fig.2C and Fig.S2C). (ii) In reporter assays, when motif 1 (GCCAT) was mutated to GCCAC the activity of M2 was diminished (Fig.1C and Fig.S1A), suggesting GCCAC is not compatible for YY1 binding.

A regulatory role of YY1 for Tf (but not Gf) expression is further supported by our secondary analysis of published YY1 ChIP-seq data (39). Importantly, apart from a small LTR subfamily, Tf_I and Tf_II are two TE subfamilies with the highest enrichment of YY1 binding in mESCs (Fig.6). None of the other evolutionarily young mouse L1 subfamilies, including Gf_I and Gf_II, were enriched of YY1. Unlike Tf_I and Tf_II monomers, Tf_III monomers are degenerate at the underlined nucleotide position in the YY1-binding site (GTCGCCATCT<u>T</u>G). This nucleotide position is highly conserved among YY1-binding sites (Fig.1B). The deviation from the consensus may lead to reduced YY1 interaction and thus explain why Tf_III subfamily is not enriched for YY1 binding in the ChIP-seq data.

Our finding that YY1 is a transcriptional activator of Tf subfamilies sheds light on their high retrotransposition activity relative to other evolutionarily young mouse L1 subfamilies. Among a compilation of 12 germline mutations caused by L1 insertions, all five full-length insertions belong to Tf_I and Tf_II subfamilies, one rearranged insertion is also a Tf element, and the remaining insertions could not be assigned to a specific subfamily due to insufficient sequence information (50). Similarly, all 11 de novo mouse L1 insertions found in a recent lineage analysis of heritable insertions are Tf elements (51). Heritable insertions occur during early embryogenesis and/or gametogenesis when L1 promoter is hypomethylated (51–53). An activating role of YY1 for Tf elements during these developmental stages is supported by preferential binding of YY1 to unmethylated motifs. In this regard, IAP exemplifies methylation-dependent regulation of YY1 function (47,54). In undifferentiated F9 cells, the YY1-binding motif in IAP promoter is unmethylated; consequently, YY1 binds to and silences IAP promoter. In differentiated F9 cells, however, the YY1-binding motif in IAP promoter becomes methylated, blocks YY1 binding, and enables IAP expression. The inability of YY1 to bind to methylated motifs has also been demonstrated in multiple imprinted genes (55,56) and, most recently, across the genome (57). Thus, we expect YY1 to behave similarly at Tf promoters: binding and activating unmethylated motifs but unable to bind and function when Tf monomers are methylated. In this context, the ubiquitously expressed YY1 functions as “a permanently present basal transcription factor whose activity is controlled by secondary events” (as proposed in (19)), such as DNA methylation.

Importantly, our observation helps to explain the synergy between Tf monomers. We previously dissected the relative contribution of M2, M1, and the tether sequence to the overall promoter activity for A_I, Gf_I, and Tf_I subfamilies (34). For A_I subfamily, M2 is the major contributor, M1 has minimal activity, and the tether negatively regulates M2 in the context of two-monomer 5’UTR. The Gf_I subfamily has the lowest promoter activity tested, with contribution from a synergistic interaction between M2 and the tether. For Tf_I subfamily, it appears that M2 and M1 are synergistic while the tether is additive to the overall two-monomer promoter activity.

However, trans-acting factors mediating the synergistic interaction between Tf_I M2 and M1 were unknown. In the current study, we showed that an intact YY1 motif was required for the activity of each monomer when tested on its own and in the context of two tandem monomers and that simultaneously mutating both YY1 motifs eliminated the two-monomer promoter activity (Fig.1 and Fig.S1). YY1 can form dimers and high-order oligomers (58). YY1 dimerization promotes interactions between enhancers and promoters (21,23). Thus, it is conceivable that tandem arrayed Tf monomers may be bridged together via YY1 binding and multimerization.

Under this model, distant monomers may act like enhancers and synergistically boost the activity of the proximal monomers (Fig.7; model). By positioning multiple YY1-binding monomers immediately upstream, Tf_I and Tf_II 5’UTRs are configured like a housekeeping gene promoter, which tends to have built-in enhancer activities mediated by GABPA or YY1 motifs (59). As a result, it may help to minimize the influence of distal enhancers on L1 promoter activity. This model is consistent with a recent report of methylation patterns across the entire mouse L1 5’UTR in undifferentiated mESCs: the inner monomers are consistently hypomethylated in elements containing three or more monomer units (60).

**Figure 7.**
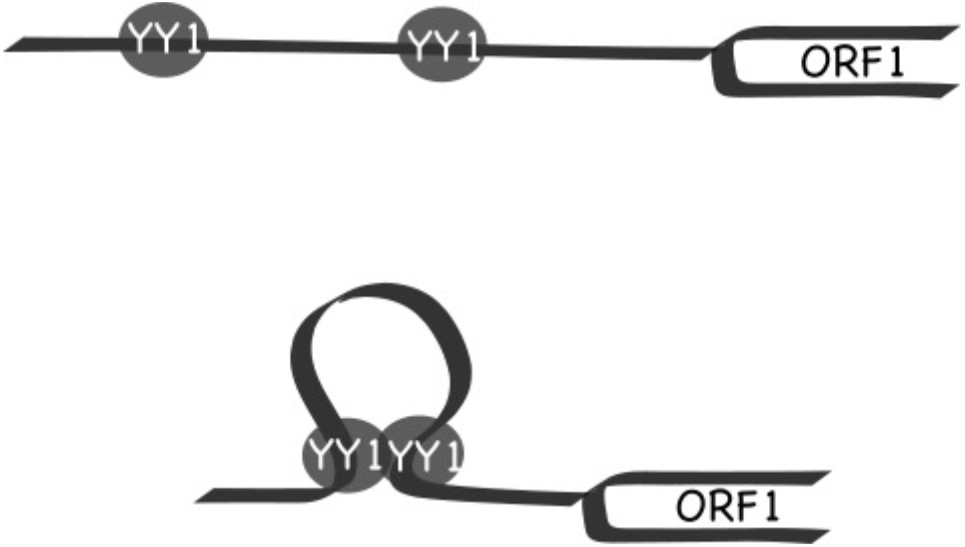
Model of YY1-mediated promoter synergy between Tf monomers. (Top) A two-monomer 5’UTR promoter sequence is positioned upstream of the ORF1 coding sequence. Each monomer harbors a functional YY1-binding site, which is occupied by a YY1 molecule. (Bottom) The two YY1 molecules dimerize and form a transcription hub, resulting in synergistic increase in promoter activity. In this model, the upstream monomer functions as an enhancer for the downstream monomer.

The large number of YY1-binding sites in TEs has implications for gene regulation at the genome level. A total of 118 subfamilies showed varied enrichment of YY1 occupancy in all three ChIP-seq libraries from mESCs (Fig.6B), including 99 subfamilies from the LTR class, 11 subfamilies from the SINE class, 7 subfamilies from the LINE class, and one subfamily from the DNA transposon class (Fig.6C). Although enriched to less extent at individual subfamily level than Tf_I and Tf_II, together these subfamilies provide a formidable collection of YY1-binding sites. There is a striking parallel in the human genome. YY1-binding sites have been identified in all four classes of human retrotransposons. For LTR element, a YY1-binding site is found in the U3 region of HERV-K and activates HERV-K transcription (61). For the SINE class, a YY1-binding site is in the left monomer of Alu, downstream from the RNA Pol III promoter (62–64). A DNA fragment containing this binding motif interacts with recombinant YY1 protein in vitro although showing lesser affinity when compared to a fragment containing the canonical YY1-binding site (63). Lastly, a composite YY1-OCT4 binding motif or a YY1 motif alone is enriched in SVAs that are transcribed in human induced pluripotent stem cells (65). Thus, in both mouse and human genomes, YY1-binding motifs in TEs may contribute to the enhancer-promoter interaction network (21). Consistent to this hypothesis, full-length Tf elements display hypomethylated monomers (60) and increased chromatin accessibility (66) in mESCs.

## Data Availability

All data generated or analyzed during this study are available from the corresponding author on reasonable request. Most, if not all, of such data are included in supplemental information.

## Supplemental Figures

Figure S1. Effect of YY1 motif mutants on Tf_I promoter activity in NIH/3T3 cells.

Figure S2. Interaction of YY1 protein from NIH/3T3 cells with motif 1 but not with a pseudo motif in Tf_I monomers.

Figure S3. Knockdown of YY1 protein and its impact on the Tf_I promoter activity in F9 cells.

Figure S4. Promoter activities of YY1 motif variants of the Gf_I subfamily in NIH/3T3 cells.

Figure S5. Lack of interaction of YY1-containing nuclear protein extract with a putative YY1 binding motif in Gf_I monomers.

Table S1. Promoters used in single vector dual luciferase assays.

Table S2. DNA fragments used in EMSA.

Table S3. Enrichment of YY1 occupancy at TE subfamilies in mESCs.

## Acknowledgements

We thank An lab members for discussion and support, and Luke Gassman for help with high performance computing.

## Authors’ contributions

KS, GIN, RN, and LK performed experiments; PY conducted computational analyses; KS and WA designed the project and wrote the manuscript; WA directed the project.

