## Supplemental Figure S1-S5 for "YY1 is a transcriptional activator of mouse LINE-1 Tf subfamily"

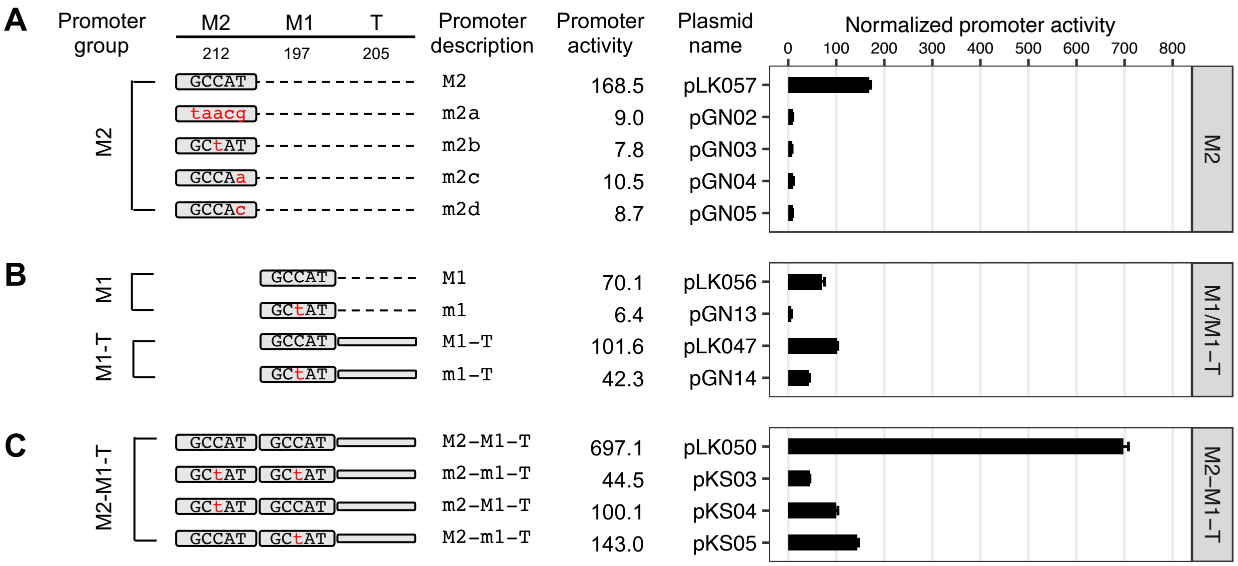


**Figure S1. Effect of YY1 motif mutants on Tf_I promoter activity in NIH/3T3 cells.** (**A**) Normalized promoter activity of M2 constructs. Mutation to consensus YY1 binding motif 1 (GCCAT) is indicated by lowercases in red. The four mutant promoter variants uniformly displayed minimal promoter activity, corresponding to 16- to 22-fold reduction as compared to the wild-type. (**B**) Normalized promoter activity of M1 and M1-Tether (M1-T) constructs. Mutant M1 (m1) showed 11-fold less activity than M1. A 2.4-fold reduction was seen in m1-T as compared to M1-T. (**C**) Normalized promoter activity of M2-M1-T constructs. The promoter activity was reduced by 16-fold when both monomers were mutated. Mutation to one monomer at a time showed activity from the other monomer and tether. For panels A-C, sequence organization of the promoters is illustrated on the left side. The length of M2, M1, and tether for each promoter is annotated (in base pairs). The dashed line represents domain(s) that were removed in reference to the two-monomer 5'UTR sequence (M2-M1-T). The x-axis indicates the normalized promoter activity, which is also listed under column "promoter activity" for each promoter variant. The positive control construct, pCH117, had a normalized promoter activity of 1526.3. Error bars represent standard errors of the mean (n = 4).

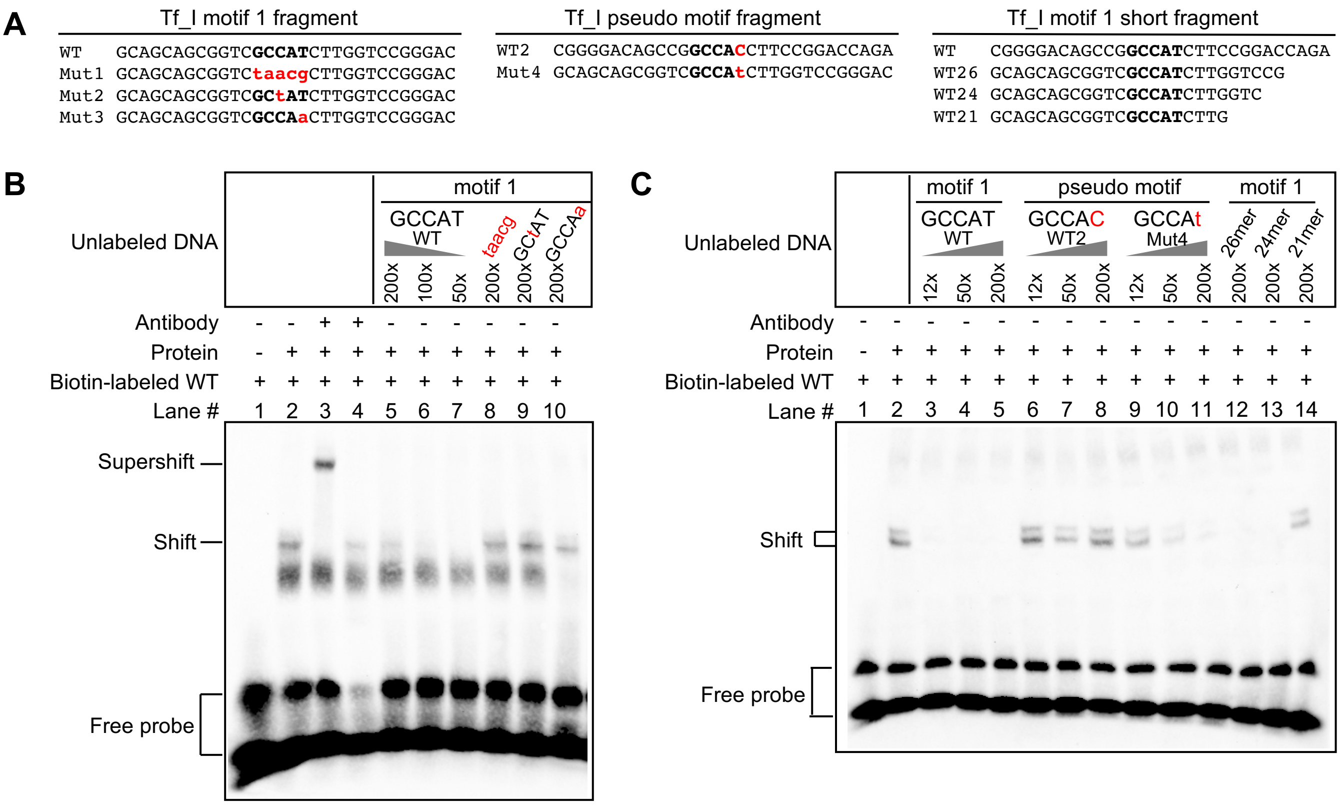


**Figure S2. Interaction of YY1 protein from NIH/3T3 cells with motif 1 but not with a pseudo motif in Tf_I monomers.** (**A**) Wild-type and mutant DNA fragments used in electrophoretic mobility shift assays (EMSA). Each fragment was formed by annealing a sense stranded oligo (shown) with the corresponding antisense oligo (not shown). Mutations in the core binding motif are indicated by lowercases in red. The wild-type pseudo motif (WT2) differs from motif 1 (WT) at the fifth position of the core binding sequence. (**B**) EMSA with Tf_I motif 1 fragments. The presence or absence of a biotin-labeled WT probe, antibody, and nuclear protein extract from NIH/3T3 cells is indicated by "+" or "-" symbols. Lanes 5-10 had unlabeled DNA fragments as competitors in molar excess as indicated. A shift was observed in the presence of nuclear protein extract. A supershift was observed with the addition of YY1 antibody (lane 3) but not by mouse IgG (lane 4). The shift was diminished by increasing amount of unlabeled WT fragment but not by unlabeled mutant DNA fragments (Mut1, Mut2, and Mut3). (**C**) EMSA with Tf_I pseudo motif fragments and shortened motif 1 fragments. The presence or absence of a biotin-labeled WT probe, antibody, and nuclear protein extract from NIH/3T3 cells is indicated by "+" or "-" symbols. Lanes 3-14 had unlabeled DNA fragments as competitors in molar excess as indicated. The shift was diminished by an unlabeled mutant fragment containing the consensus core motif (Mut4; lanes 9-11) by not by unlabeled wild-type pseudo motif fragment (WT2; lanes 6-8). Shortened motif 1 containing DNA fragments (26bp or 24bp) were able to inhibit the shift but 21bp fragments were not as effective (lane 14).

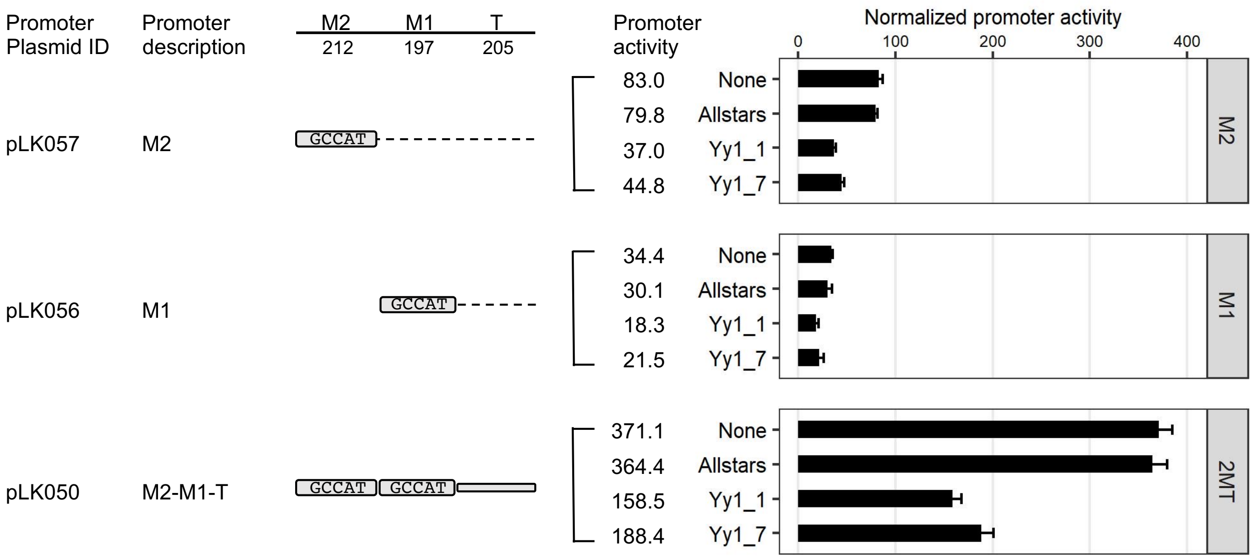


**Figure S3. Knockdown of YY1 protein and its impact on the Tf_I promoter activity in F9 cells.** Normalized promoter activity for three Tf_I promoter constructs (M2, M1, and M2-M1-T) under siRNA knockdown. For each promoter variant, cells were cotransfected with the promoter construct and with or without a siRNA (y axis; none = no siRNA). In reference to cells treated with Allstars, Yy1_1 siRNA treated cells showed 46.3%, 60.7%, and 43.5% of the activity for M2, M1, and M2-M1-T, respectively. In comparison, Yy1_7 treated cells showed 56.1%, 71.4%, and 51.7% of the activity for M2, M1, and M2-M1-T (also marked as 2MT), respectively. The positive control construct, pCH117, had a normalized promoter activity of 950.2. Error bars represent standard errors of the mean (n = 4).

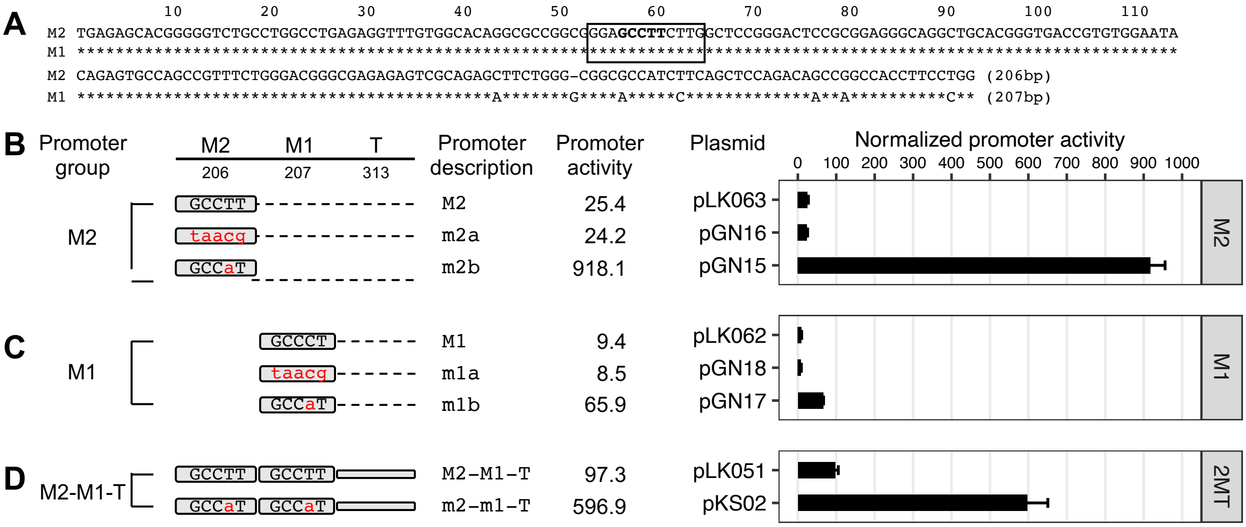


**Figure S4. Promoter activities of YY1 motif variants of the Gf_I subfamily in NIH/3T3 cells.** (**A**) Alignment of Gf_I monomer 2 (M2) and monomer 1 (M1) consensus sequences. In the M1 sequence, nucleotide positions identical to M2 are marked by asterisks. Sequence gaps are represented by dashes. A previously predicted YY1 binding motif is located between nts 53-64 (solid box, termed "Gf_I motif"). Promoter activity is assessed using Dual-luciferase reporter assay. **(B)** Normalized promoter activity of monomer2 (M2) constructs. Mutation to Gf_I motif (GCCCT) is indicated by lowercases in red. The m2a showed minimal change in promoter activity. However, changing to the consensus YY1 motif (m2b) elevated the M2 promoter activity by 36.1-fold. **(C)** Normalized promoter activity of monomer1 (M1) constructs. The mutant monomer 1 (m1a) showed minimal change in promoter activity. Changing to the consensus (m1b) showed 7.0 times higher signal compared to M1. **(D)** Normalized promoter activity of monomer2-monomer1-Tether (M2-M1-T; or 2MT) constructs. A 6.1-fold higher activity was observed upon changing both Gf_I motifs to the consensus sequence. The positive control construct, pCH117, had a normalized promoter activity of 1942.5. Error bars represent standard errors of the mean (n = 4).

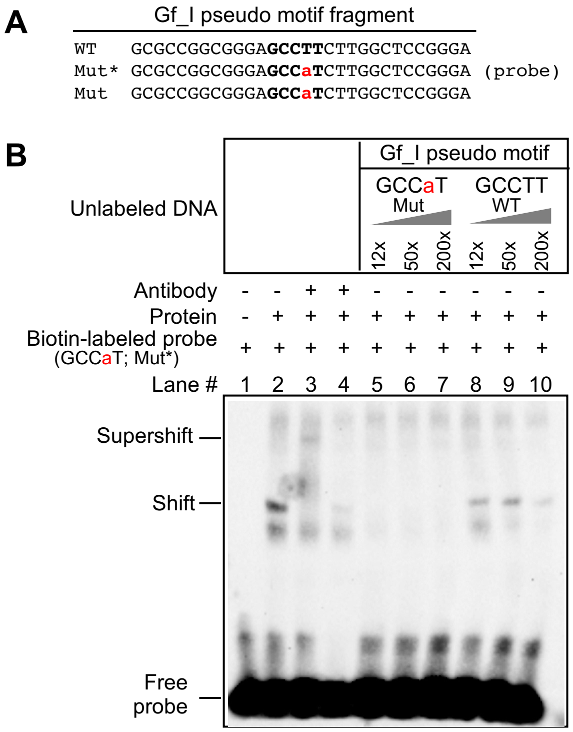


**Figure S5. Lack of interaction of YY1-containing nuclear protein extract with a putative YY1 binding motif in Gf_I monomers.** (**A**) Wild-type and mutant DNA fragments used in electrophoretic mobility shift assays (EMSA). Each fragment was formed by annealing a sense stranded oligo (shown) with the corresponding antisense oligo (not shown). Mutation in the core binding motif is indicated by a lowercase letter in red. In the mutant variant (Mut), the wild-type pseudo motif (WT) was mutated at the fourth position to restore the consensus core binding sequence. (**B**) EMSA with Gf_I pseudo motif fragments. The presence or absence of a biotin-labeled Mut probe (GCCaT), antibody, and nuclear protein extract from NIH/3T3 cells is indicated by “+” or “-” symbols. Lane 3 had YY1-specific antibody and lane 4 had mouse IgG as a control. Lanes 5-10 had unlabeled DNA fragments as competitors in molar excess as indicated.
